# Isolation and characterization of novel *Klebsiella* phages from Benin and their antibiofilm activities on multidrug resistant and hypervirulent strains of *Klebsiella pneumoniae*

**DOI:** 10.64898/2026.07.25.740742

**Authors:** Alidehou Jerrold Agbankpe, Kafayath Fabiyi, Yaovi M. Gildas Houumanou, Gracia Ruth Nougbologni, Roubaya Balarabe, Slawomir Michniewski, Jules Beau-Gard Hougbenou, Esther Deguenon, Victorien Dougnon, Honoré Bankolé, Lamine Baba-Moussa, Rashid Nazir, Martha R.J. Clokie

## Abstract

Hypervirulent *Klebsiella pneumoniae* represents a growing clinical threat, particularly where multidrug resistance limits treatment options. Here, we report the isolation and characterization of three novel lytic phages from Benin (Kp1Bj_HH11_M23, Kp2Bj_LN294_M23, and Kp10Bj_LN54_14). These Myoviruses, belonging to the genera *Marfavirus* and *Jiaodavirus*, display broad host range activity against multidrug-resistant and hypervirulent *K. pneumoniae* strains. Genomic analysis confirmed the absence of virulence and antimicrobial resistance genes. The phages exhibit rapid adsorption, short latency periods, and high burst sizes (119–2208 PFU/cell). All three phages significantly inhibited biofilm formation and reduced established biofilms in vitro. Their stability across a wide range of temperatures and pH further supports their potential for therapeutic development. Together, these data highlight the value of locally sourced phages as candidates for tackling region-specific antimicrobial resistance challenges.

## 1 Introduction

Antibiotics have been essential as a first line of defence against pathogens, playing a crucial role in the fields of human and animal health (Manyi-Loh et al., 2018; Rahman et al., 2025). However, several decades since the introduction of antibiotics, the world is facing an uncertain situation in the advent of the post-antibiotic era. Basically, the uncontrolled use of antibiotics in human and veterinary medicine has amplified and accelerated the evolution of antibiotic resistance (Schooley, 2017). The problem of transmission and dissemination of resistance has been intensified by environmental contamination of resistant bacteria and resistance genes from people and animals (Carvalho et al., 2017). To this end, communities and hospitals feel invaded by the increasing spread of antibiotic-resistance bacteria is threatening our ability to treat serious infections and represents a significant public health problem globally (Salam et al., 2023). Gram-negative bacteria constitute a significant fraction of multidrug resistant pathogens, including *Klebsiella pneumoniae*, for which there is an urgent need to research and develop new treatments (*WHO*, 2024).

*K. pneumoniae* infections are often considered one of the deadliest infections for infants, the elderly, and immuno-compromised patients. Despite the development of broad-spectrum antimicrobials, more than 50% of mortality rates have been recorded in individuals infected with *K. pneumoniae* (Bassetti et al., 2018; Fage et al., 2021). Antimicrobial resistance (AMR) in *K. pneumoniae* is a major clinical challenge, particularly with the emergence of strains combining multidrug resistance and hypervirulence (Arcari et al., 2023; Gan et al., 2025). In early 2024, the World Health Organization (WHO) reported the global emergence of *K. pneumoniae* ST23 strains carrying carbapenemase genes, which confer resistance to last-resort antibiotics (Abdel-Rahman et al., 2026).

This coexistence of hypervirulence and multidrug resistance is associated with severe infections, high mortality, and limited treatment options, particularly in low- and middle-income countries (LMICs) (Kansal et al., 2026). This issue has led to the emergence of phage therapy as a promising alternative. Despite the public health burden that hypervirulent and multidrug resistant *K. pneumoniae* can pose in LMICs, there is a lack of region-specific data on phages that target clinically relevant strains of *K. pneumoniae* in Africa. To address this gap, we have undertaken to isolate and perform molecular characterization of phages from Benin that target hypervirulent and multidrug resistant strains of *K. pneumoniae*, both to analyze the genetic diversity of these phages and to explore their potential as therapeutic agents.

## 2 Material and methods

### 2.1 Bacterial strains

A collection of sixty clinical strains of hypervirulent and multidrug resistant *K. pneumoniae* were used in this study. The strains were obtained from the bacterial strain collections of the Becky Meyer Centre for Phage Research, University of Leicester, United Kingdom (UK) and the Research Unit in Applied Microbiology and Pharmacology of natural substances, University of Abomey-Calavi, Benin. These strains were isolated from clinical samples of urine (midstream or catheter specimen urine), blood, rectal swab, sputum and endotracheal secretions, and with a diversity of capsular loci (KL) (Supplementary Table S1). All *K. pneumoniae* strains were grown at 37°C in either Luria-Bertani broth (LB-Thermo Fisher Scientific, UK) at 200 rpm or on LB 1.5% (w/v) agar plates and then used as hosts for phage isolation and host range evaluation.

### 2.2 Phage isolation, purification and visualisation

Phages were isolated from hospital effluent, farmer’s stools and pig droppings collected from pig farms located on Lake Nokoué in Benin (Supplementary Table S2) using *K. pneumoniae* host strains as previously described with slight modification (Haines et al., 2021). Samples (50 mL) were centrifuged at 4,200 x g for 10 min to remove debris and other undissolved particles, and the supernatant was filtered using a 0.45µm filter. Filtrate (10 mL) was enriched by bacterial culture (100 µl) in LB broth (10 mL) supplemented with 40 µl CaCl2 (1M) and incubated overnight at 37°C with shaking (200 rpm). The mixed culture solution was centrifuged at 4,200 x g for 15 min, and the supernatant was filtered using a 0.22 µm filter. Then, enriched filtrate (100 µl), bacterial culture (100 µl) and LB 0.6% (w/v) agar (3 mL) were poured onto a LB 1.5% (w/v) agar plate (90 mm petri dishes) and incubated overnight at 37°C. Single plaques were picked and transferred to SM Buffer (500 µl). The mixture was shaken for 1h by inversion, centrifuged at 4,200 x g for 30 min and was filtered using a 0.22 µm filter.

Transmission electron microscopy (TEM) imaging for the phages was performed at a microscopy core facility of the University of Leicester, UK. 5µL of phages were negatively stained with 1% (w/v) uranyl acetate on 3 mm carbon-coated copper grids and visualised with a JEM-1400 transmission electron microscope (JEOL UK Ltd., UK) with an accelerating voltage of 120kV. Digital images were collected using a Xarosa digital camera (EMSIS, Germany) with Radius software.

### 2.3 Titration, host range and Efficiency of Plaquing (EOP)

The titration of each phage was performed using the method previously described by El-Telbany et al. (2021). A serial dilution of each phage was performed in SM buffer (900 µL). Bacterial culture (250 µL, OD_600_ of ∼0.3) was added to 8 ml LB 0.6% (w/v) agar and poured onto LB 1.5% (w/c) agar plates (120 x 120 mm petri dishes) for bacteria lawn. Then, each dilution (10 µL) was spotted on LB agar plates and incubated at 37°C overnight. The titer in plate-forming units per millilitre (PFU/mL) of each phage was calculated using the following formula: PFU/mL = (number of plaques x dilution factor)/volume spotted (mL).

Host range analysis was performed using a panel of fifty-eight *K. pneumoniae* strains (Supplementary Table S1) by spot test. Phage stock (10 µL) was spotted onto the plate (bacterial lawn) and incubated overnight at 37°C. The plaques were graded to complete lysis and hazy lysis. Moreover, susceptibility of the tested strains to the phages was also performed using the efficiency of plaquing (EOP) method (El-Telbany et al. 2021; Kutter, 2009). Serial dilutions were made for each phage and 10 µL of the dilutions were spotted on lawns. Plaques were counted, and EOP was calculated using the following formula: average PFU/mL on tested strain/average PFU/mL on potential host.

### 2.4 Phage DNA extraction and sequencing

Phage DNA extraction was performed using the phenol-chloroform-isoamyl alcohol (PCI) method (Nale et al., 2015). Phage lysate (1mL) was treated with 12.5 μL of 1M MgCl_2_, 10 µL of DNase I (30 mg/ml) and 1 µL of RNase A (100 mg/mL), followed by incubation for 30 min at room temperature. Proteins were removed and phage DNA was purified by two cycles of PCI extractions. Then, DNA was precipitated with 95% ethanol and 3M sodium acetate solution. After washing in 70% ethanol, the pellets were resuspended in 50 µL of water. DNA concentrations were quantified using a Qubit 2.0 fluorometer. Illumina sequencing libraries were prepared using the tagmentation-based and PCR-based Illumina DNA Prep kit and custom IDT 10bp unique dual indices (UDI) with a target insert size of 320 bp. No additional DNA fragmentation or size selection steps were performed. Illumina sequencing was performed on an Illumina NovaSeq 6000 sequencer for 151bp paired-end reads. Demultiplexing, quality control and adapter trimming was performed with bcl-convert (v4.1.5).

### 2.5 Bioinformatics analysis

#### 2.5.1 Phage Genome Identification and Initial Curation

Raw reads of phages were assembled with Spades v4.2.0. and added to a curated collection of 39 publicly available *K. pneumoniae* phage genomes. For each sample, assembled contigs in FASTA format were quality controlled with CheckV v1.0.3 and analyzed using VirSorter2 (v2.2.3), focusing on identification of dsDNA phages, with the dsDNAphage group enabled and a minimum contig length threshold of 1,500 bp. All resulting outputs were organized in structured directories for batch processing. To supplement VirSorter2 predictions and identify additional phage sequences, a complementary BLASTn-based filtering strategy was applied. All contigs were searched against the NCBI nucleotide (nt) database using BLASTn, with filters set to ≥75% identity, ≥25% query coverage, and E-value ≤1e-5. Resulting hits were retained if their annotations included the terms “phage”, “virus”, or “bacteriophage”. Corresponding contig headers were extracted using seqtk to generate refined FASTA files representing likely phage genomes.

#### 2.5.2 High-Confidence dsDNA Phage Sequence Selection

To increase the specificity of phage detection, candidate sequences were filtered for high-confidence double-stranded DNA (dsDNA) phages based on classification results from VirSorter2. Only contigs classified under the dsDNA phage group with a final_max_score of 1.0 were retained. Contig identifiers were mapped to headers in the final-viral-combined.fa files to extract full sequence entries. These sequences were then compiled into sample-specific FASTA files as well as individual contig files (*.fa) using a custom extraction pipeline. Curated phage sequences were consolidated across all samples into a central directory for downstream analyses. Empty files and low-quality outputs were removed to ensure a clean dataset for further processing.

#### 2.5.3 Phage Genome Quality Assessment

All candidate phage genomes were evaluated using CheckV (v1.0.1) to assess genome completeness, contamination, and provirus content, using the CheckV database (v1.5). Quality summaries were parsed and centralized, and provirus sequences (viruses.fna) were extracted and renamed for standardized downstream analysis. Quast was then used to generate the overall assembly statistics on the final phage genomes retained for analysis.

#### 2.5.4 Phage Genome Annotation

High-quality phage genomes were annotated using two complementary pipelines:

- Cenote-Taker3 was applied for structural and functional annotation, with a focus on viral hallmark genes including those encoding capsid and tail proteins.
- Pharokka (v1.7.1) was used to annotate protein-coding genes across phage genomes based on the PHROGs and CARD databases. Annotation was performed on each phage genome individually, and results were stored in organized output directories. Visual summaries of the gene content were generated using pharokka_multiplotter.py.

#### 2.5.5 Taxonomic Classification of Phage Genomes

To confirm and refine the taxonomic identities of the phage sequences, Kraken2 (v2.1.2) was used to classify all curated genomes against two viral databases. One Kraken2 run utilized a custom viral/phage-specific Kraken2 database, while another employed a standard RefSeq Kraken2 database. Classification reports were parsed to assess taxonomic diversity and ensure inclusion of known phage lineages.

#### 2.5.6 Phylogenetic Analysis of Phage Genomes

To investigate the evolutionary relationships among phage genomes, we constructed phylogeny based on single nucleotide polymorphisms (SNPs) in accessory genome content. This analysis included the three reference genomes (from this study) and thirty-nine high-quality *K. pneumoniae* phage genomes downloaded from NCBI as publicly available genomes (Supplementary table S3). We employed a pangenomic approach to explore gene content diversity among all forty-two phage genomes. Each genome was annotated using Pharokka, a tool specifically designed for phage genome annotation. The resulting GFF3 files were used as input for Roary, which clustered orthologous genes and produced a gene presence/absence matrix. A phylogenetic tree was subsequently inferred based on this binary gene content matrix. This approach allowed us to investigate evolutionary relationships shaped by gene gain and loss, capturing genomic variability not evident through SNP analysis alone and providing a broader view of the diversity and relatedness among phage genomes. Average Nucleotide Identity (ANI) analysis was performed between the three phages using fastANI V1.34.

### 2.6 Optimal multiplicity of infection (MOI)

The MOI for each phage was determined using the method previously described by Hans et al. (2023). Bacterial culture (150 µL) was mixed with different dilutions of phage suspension at MOIs of 0.001, 0.01, 0.1, 1, 10, and 100. The mixture was added to 5 ml of LB and incubated at 37°C, 200 rpm for 18h. The culture was centrifuged at 4,200 x g for 10 min, and the supernatant was filtered using 0.22 µm filter. Then, phage titer under different MOI was determined using double layer agar method (Haines et al., 2021), and the highest phage titer was corresponded to the optimal MOI.

### 2.7 Adsorption assays and one-step growth curve

Adsorption assays were performed as previously described with some modification (Storms and Sauvageau, 2014). Bacterial culture (900 µL) was aliquoted to Eppendorf tubes labelled 0; 5; 10; 15; 20; 25; 30; and 60 min. Then, phage (100 µL) was added to each Eppendorf tubes at an optimal MOI and incubated at 37°C. At each time point, mixture was removed and centrifuged at 12,000 rpm for 5 min, and the supernatant was filtered using 0.22µm. Serial dilution of filtrate was carried out, and the titer of unadsorbed phage was determined using the double layer agar method.

One-step growth was carried out to understand the growth pattern of *Klebsiella* phage from Benin. Phage was added to bacterial culture at an optimal MOI. The mixture was incubated at 4°C for 20 min to allow sufficient adsorption of phage to the host, and was centrifuged at 4,200 rpm for 10 min. Then, pellet (bacteria) were collected. Pellet was washed twice with SM buffer and resuspended in LB (10 mL). This suspension (1 mL) was aliquoted to Eppendorf tubes labelled 0; 5; 10; 15; 20; 30; 60; 90; and 120 min and was incubated at 37°C, 200 rpm. At each time point, the mixture was removed and centrifuged at 12,000 rpm for 5 min and was filtered. Serial dilution of filtrate was carried out, and the titer of phage was determined using the double layer agar method. The number of viral particles produced from the infection of a single cell, called the burst size, was calculated with titer at latent phase/titer at the plateau (Zhao et al., 2019).

### 2.8 Antibiofilm activities of phages against *K. pneumoniae* host strains

*K. pneumoniae* produces a capsular polysaccharide that can be released into the extracellular environment to form biofilms that can enhance host defence and tolerance against antimicrobials (Anderl et al., 2000). Planktonic cells killing was performed and the inhibition of biofilm formation of *K. pneumoniae* strains by phages isolates for 48 hours as well. The experiment was carried out over 4 days. Day 0: Four 12-well plates were labelled P1, P2, P3 and P4 respectively. Bacterial culture (1 mL, OD_600_ of ∼0.3) was added to wells A1-2, B1-2 and C1-2. Then, LB broth (1 mL) was added to wells A4, B4 and C4, which served as negative controls. The four plates were incubated at 37°C, 100 rpm for 24 h. Day 1: planktonic cultures (500 µL) were removed from plates P3 and P4, taking care to not disturb the biofilms. Each planktonic culture was replaced by LB broth (500 µL). Both plates were incubated again at 37°C, 100 rpm for 24 h. All the planktonic cultures were removed from plates P1 and P2 and the wells were washed 3 times by gently adding and removing PBS (1 mL). Then, phage (500 µL) was added to wells A1, B1 and C1 (phage treated wells), and LB broth (500 µL) were added to wells A2, B2 and C2 (bacterial control), and 500 µL of LB broth to wells A4, B4 and C4 as well. Plates were incubated at 37°C, 100 rpm for a further 24 h. Day 2: on plate P1, planktonic cultures (500 µL) was removed from each of the phage treated and bacterial control wells and put into Eppendorf tubes and was washed three times by centrifuging at 12,000 rpm for 15 min and resuspending with PBS (500 µL). The final pellet was resuspended with 1 ml of PBS and serial dilution was conducted and bacteria was counted (CFU/ml) by spotting the dilution on to LB 1.5% (w/c) agar. The plates were incubated at 37°C for 24 h. For biofilms, the wells were washed three times by gently adding and aspirating PBS (1 mL). Then PBS was added, and the bottom of wells was scraped to resuspend the biofilm. Serial dilution of the biofilm culture was performed, and bacteria was counted (CFU/ml). For the P2 plate, the plankton cultures were aspirated, and the wells were washed three times by adding and gently aspirating PBS. Then, 500 µL of 0.1% filtered crystal violet (CV) was added to the wells and placed on a rocker for 15 min at room temperature. The wells were gently washed under the tap to remove any unbound CV. Ethanol (100%, 1 mL) was added, and the plate was incubated at room temperature for 10 min. The well content (200 µL) was transferred to a new 96-well plate and the OD_600nm_ was measured using the BMG Labtech SPECTROstar Omega (Sarstedt, Germany). Then, experiments carried out on plates P1, and P2 on Day 1 were repeated on plates P3 and P4. Day 3: The experiments carried out on plates P1, and P2 on Day 2 were repeated on plates P3 and P4.

### 2.9 Thermal and pH stability

In Eppendorf tubes labelled 4, 10, 20, 30, 40, 50, 60, 70 and 80°C, phages (100 µL) were added to SM buffer (900 µL). Tubes identified as 4 and 10°C were incubated in the refrigerator and those at 20, 30, 40, 50, 60, 70 and 80°C were incubated in thermomixers for 1h. Then, tubes were brought to room temperature for 1 h on the bench, and serial dilution of each tube (100 µL) was carried out. Titer of phage was determined using the double layer agar method. For pH stability, SM buffer was adjusted by adding HCl or NaOH to obtain the desired pH using an automatic pH meter.

### 2.10 Ethical approval

The study was approved by the National Health Research Ethics Committee of Benin (N◦67/MS/DC/SGM/DRFMT/CNERS/SA of 20th April 2021) and farmers provided written informed consent.

## 3 Results

### 3.1 Isolation and morphology of phages

The three phages were isolated from hospital effluent (Kp1Bj_HH11_M23), farmers’ stools (Kp10Bj_LN54_14) and pig feces (Kp2Bj_LN294_M23) on multidrug resistant and hypervirulent *K. pneumoniae* (Supplementary Table S2). The phages Kp1Bj_HH11_M23 and Kp2Bj_LN294_M23 formed round 1±1 mm translucent plaques on *K. pneumoniae* ST48 lawns (Figures 1A and 1B respectively). Kp10Bj_LN54_14 plaques had diameter of 2±1 mm on *K. pneumoniae* (unknown ST) lawns (Figures 1C). Transmission electron microscopy (TEM) analysis of phages particles indicated that Kp1Bj_HH11_M23, Kp2Bj_LN294_M23, and Kp10Bj_LN54_14 phages had an icosahedral heads with a diameter of 86.67±3.15, 93.33±2.55, and 93.33±3.10 nm respectively with long contractile tails length of 126.67±9.55, 126.67±6.10, and 126±10.06 nm respectively (Figures 2A, 2B, and 2C).

**Figure 1.**
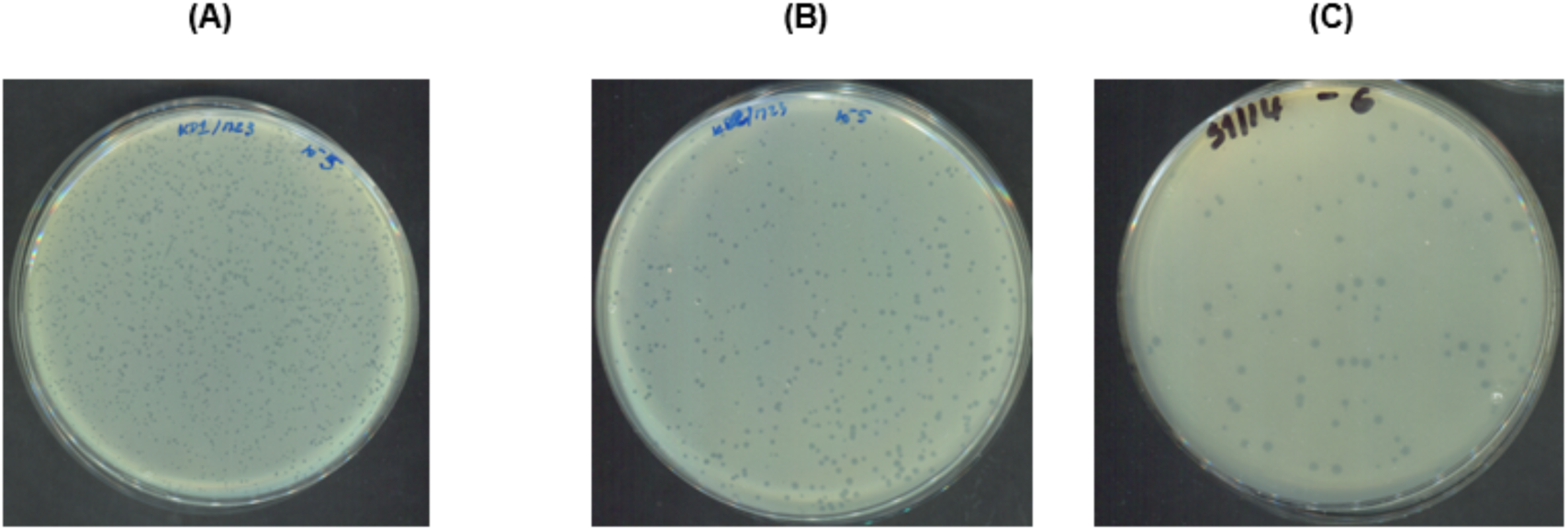
Plaque morphology of three phages formed in a double-layer agar plate with K. pneumoniae host strains (supplementary Table S1). **(A)** Plaque morphology of phage Kp1Bj_HH11_M23. **(B)** Plaque morphology of phage Kp2Bj_LN294_M23. **(C)** Plaque morphology of phage Kp10Bj_LN54_14.

**Figure 2.**
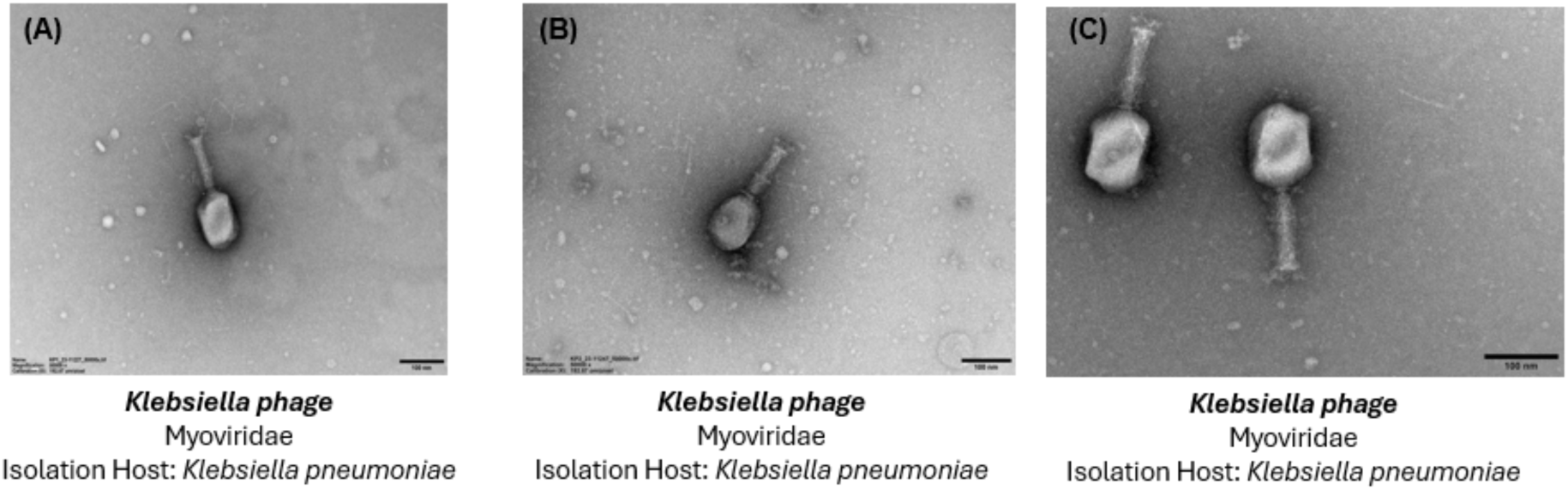
Transmission electron micrographs of phages negatively stained with 1% (w/v) uranyl acetate. The scale bar represents 100 nm. **(A)** micrograph of phage Kp1Bj_HH11_M23. **(B)** micrograph of phage Kp2Bj_LN294_M23. **(C)** micrograph of phage Kp10Bj_LN54_14.

### 3.2 Titer, host range and Efficiency of Plating (EOP) of isolated phages

The phage titre ranged from 7.10^9^ to 2.10^10^ PFU/ml. The host range was determined on a panel of 58 *K. pneumoniae* (supplementary Table S1) and showed that Kp2Bj_LN294_M23 exhibited lytic activity on 74.14% (43 out of 58) of *K. pneumoniae* strains tested with 60.47% of complete lysis plaque and 39.53% of hazy plaques. Kp1Bj_HH11_M23 and Kp10Bj_LN54_14 had lytic activities against 70.69% (41) and 68.97% (40) of *K. pneumoniae* strains tested respectively. Kp1Bj_HH11_M23 had complete lysed plaques on 60.98% strains and 39.02% had hazy lysis. 72.5% of complete lysis plaque and 27.5% of hazy lysis plaque reflects lytic activity of Kp10Bj_LN54_14. Kp2Bj_LN294_M23 phage had the largest percentage of high EOP (7.14%), followed by Kp1Bj_HH11_M23 (5%) (Table 1). The three phages exhibited lytic activity against a wide range of *K. pneumoniae* with a high diversity in capsular loci (KL114, KL53, KL8, KL62, KL64, KL124, KL102, KL57, KL39, KL48, KL17, KL20, KL2, KL30, KL51, KL105, KL146, KL28, KL122, KL38, KL3, KL163).

**Table 1.** Host range analysis and EOP of three phages against fifty-eight *K. pneumoniae* strains.

| Strains | Phages - Titer (PFU/mL) – EOP<sup>a</sup> |  |  |
| --- | --- | --- | --- |
|  | *Kp1Bj_HH11_M23* | *Kp2Bj_LN294_M23* | *Kp10Bj_LN54_14* |
| | $7.10^9$ | $2.10^{10}$ | $7.10^9$ |
| KR312 |  |  |  |
| KR358 |  |  |  |
| KR431 |  |  |  |
| KR437 |  |  |  |
| L02 | 1E-3 | 2.86E-5 | 2.86E-4 |
| L05 | 2E-2 ** | 7.5E-6 | 2.86E-5 |
| L06 |  |  |  |
| L10 |  |  |  |
| L11 | 2.1E-2 ** | 1E-2 * | 7.14E-5 |
| L13 | 7.1E-2 ** | 6E-4 | 7.14E-7 |
| L15 | 4E-4 | 1E-4 | 2.86E-5 |
| L19 |  |  |  |
| L21 | 2.86E-3 | 1.5E-3 | 7.14E-6 |
| L30 |  | 2.5E-1 ** | 2.86E-6 |
| L32 |  |  |  |
| M31 | 1.43E-5 | 1E-4 | 1.43E-6 |
| M19 | 1.43E-5 | 2.5E-4 | 1.43E-6 |
| M23 | - | - | 1.43E-4 |
| M20 | 3.6E-6 | 1E-3 | 1.43E-5 |
| M34 |  |  | 8.6E-3 |
| M36 | 1E-1 ** | 2.5E-3 | 1.43E-3 |
| M39 |  |  | 1.43E-3 |
| M40 | 5.7E-2 | 4E-4 | 1.43E-3 |
| M41 | 1.43E-4 | 1.5E-3 | 1.43E-3 |
| M54 | 1.43E-4 | 6.5E-4 | 1.43E-3 |
| M61 | 1.43E-3 | 6.75E-4 | 1.43E-3 |
| M63 | 1.43E-5 | 3.5E-4 | 7.14E-4 |
| M71 | 1.43E-5 | 1.5E-3 | 7.14E-6 |
| M45 | 1.5E-4 | 2E-3 |  |
| M49 | 7.14E-2 | 0.5*** | 0.000714 |
| M57 | 5.71E-1*** | 2E-1** |  |
| M13 | 1.43E-3 | 3E-4 | 1.43E-3 |
| M47 | 2.86E-1** | 1E-1** |  |
| M53 | 2.86E-1** | 1E-1** | 1.43E-5 |
| M11 | 1.28E-1** | 1E-1** | 2E-2** |
| M17 | 1.43E-4 | 4.75E-3 | 1.43E-3 |
| M68 | 1E-1** | 1.3E-3 | 4.3E-3 |
| M74 | 2.86E-5 | 1.1E-5 | 2.86E-5 |
| M25 | 1.5E-4 | 1E-4 | 7.14E-5 |
| M28 | 1.43E-4 | 2E-5 | 2.86E-1** |
| M48 | 5.7E-3 | 5.E-4 |  |
| M32 | 1E-3 | 6E-5 | 5.7E-3 |
| M33 |  |  | 4E-3 |
| M37 | 1E-1** | 5E-1*** | 2.86E-5 |
| M38 | 1E-5 | 3.3E-5 | 7.14E-6 |
| M42 |  |  |  |
| M52 | 1E-1** | 5E-2** | 1.43E-4 |
| M58 | 1E-4 | 1.5E-4 | 2.86E-6 |
| M65 | 2.86E-5 | 1E-5 | 1.14E-1** |
| M06 | 2.86*** | 4E-1** | 7.14E-6 |
| M35 |  | 3.5E-1** |  |
| M03 | 2.9E-1** | 50*** | 4E-2** |
| M24 | 4.3E-1** | 1E-1** | 1E-1** |
| M69 | 1E-5 | 1.43E-5 |  |
| M44 |  |  |  |
| M51 | 2.86E-6 | 1.5E-4 | 1.43E-4 |
| M67 |  |  |  |
| M75 |  |  |  |

*Host range: red colour = no visible plaque; green colour = complete lysis plaque; orange colour = hazy lysis plaque. <sup>a</sup> the numbers in the cells represent the EOP of each phage on the different K. pneumoniae strains tested. \*\*\* high EOP ( $EOP \geq 0.5$ ); \*\* moderate EOP ( $0.01 < EOP < 0.5$ ); \* low EOP ( $EOP \leq 0.01$ ). - no EOP calculation for initial host.*

### 3.3 Genomic comparison and phylogenetic analysis

The genomic sequences of Kp1Bj_HH11_M23, Kp2Bj_LN294_M23 and Kp10Bj_LN54_14 are available in the GenBank database under accession numbers PV784030, PV784031 and PV784032, respectively. Kp1Bj_HH11_M23 and Kp2Bj_LN294_M23 are *Marfavirus*, and Kp10Bj_LN54_14 is *Jiaodavirus*. Their genomes have approximately 170 kbp, 40% of GC content, and contained more than 200 viral genes. There were no genes encoding virulence factors or antimicrobial resistance (Table 2). The phylogenetic analysis showed that Kp1Bj_HH11_M23 and Kp2Bj_LN294_M23 are genetically related phages (with more than 99% ANI identity) compared to Kp10Bj_LN54_14. The phylogenetic tree based on the complete genome confirmed that all three phages belong to the *Myoviridae* family (Figure 3). Kp1Bj_HH11_M23 and Kp2Bj_LN294_M23 genomes encode 122 protein-coding genes (CDS) respectively, and Kp10Bj_LN54_14 encodes 123 CDS. A comparison of the annotated CDSs in the three phage genomes revealed that 186 CDSs in Kp1Bj_HH11_M23, 182 CDSs in Kp2Bj_LN294_M23, and 172 CDSs in Kp10Bj_LN54_14 are predicted to be hypothetical proteins or proteins of unknown function. Genomes contain known proteins related to phage tail, head, and packaging, proteins related to DNA metabolism, and host lysis proteins. Circular maps of the annotated genomes of Kp1Bj_HH11_M23, Kp2Bj_LN294_M23, and Kp10Bj_LN54_14 are shown in Figure 4.

**Figure 3.**
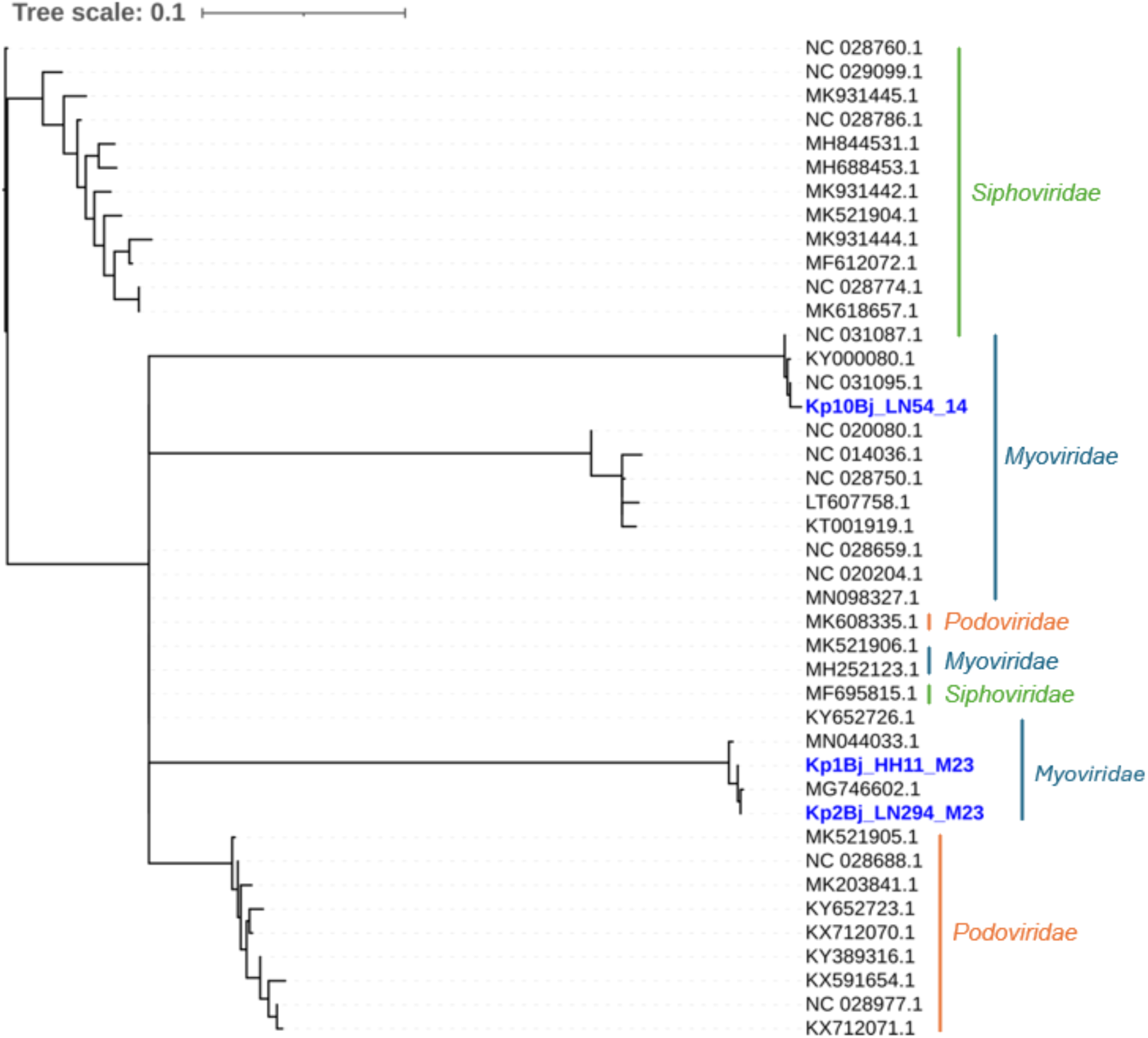
Phylogenetic tree of 42 phage genomes based on accessory genome content. Klebsiella pneumoniae reference phages are highlighted in blue.

**Figure 4.**
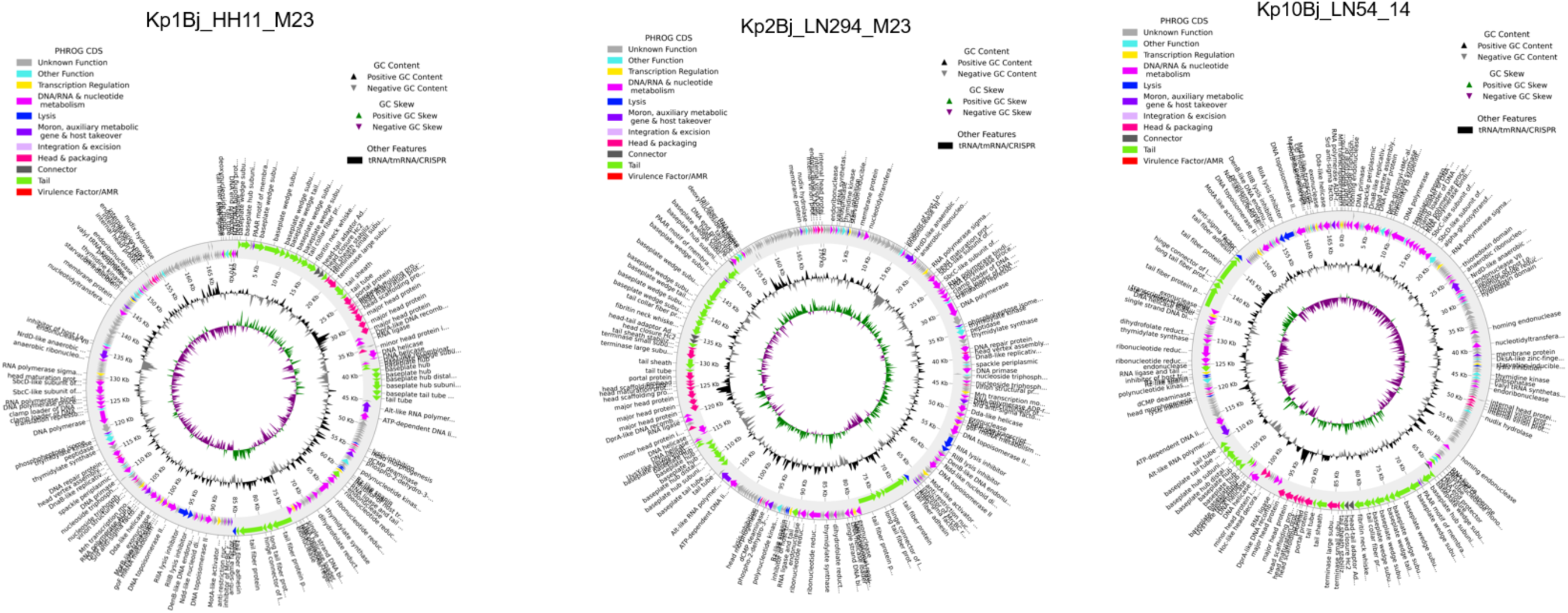
Representative genome content annotation of phages

**Table 2.** Overview of the phage genomes.

| Assembly | Kp1Bj_HH11_M23 | Kp2Bj_LN294_M23 | Kp10Bj_LN54_14 |
| --- | --- | --- | --- |
| Accession number | PV784030 | PV784031 | PV784032 |
| Family | *Myoviridae* | *Myoviridae* | *Myoviridae* |

| Genus | *Marfavirus* | *Marfavirus* | *Jiaodavirus* |
| --- | --- | --- | --- |
| Completeness | 99.34 | 99.34 | 99.07 |
| Contamination | 0.0 | 0.0 | 0.0 |
| Viral_genes | 229 | 228 | 241 |
| Number of contigs | 1 | 1 | 1 |
| Largest contig | 170568 | 170567 | 167615 |
| Total length | 170568 | 170567 | 167615 |
| GC (%) | 40.69 | 40.71 | 39.53 |
| N50 | 170568 | 170567 | 167615 |
| Antibiotic resistance genes | None | None | None |
| Identity to Kp1Bj_HH11_M23 | 100% | 99.69% | 81.44% |
| Identity to Kp2Bj_LN294_M23 | 99.70% | 100% | 80.31% |
| Identity to Kp10Bj_LN54_14 | 82.74% | 83.24% | 100% |

### 3.4 Characteristics of phages growth

Phage Kp1Bj_HH11_M23 had a higher titer at multiplicity of infection (MOI) 10, which was different from the titers obtained at the other MOIs. This difference was not significant at MOI 100, it was highly significant at MOIs 0.001, 0.01, 0.1 and 1 (Figure 5A). The highest titers of phages Kp2Bj_LN294_M23 and Kp10Bj_LN54_14 were obtained at MOIs 0.1 and 0.001 respectively. The difference between these titers and those obtained at the other MOIs (0.01, 1, 10, 100) was not significant (Figures 5B and 5C). The different kinetic adsorption curves of phages at their optimal MOI showed that Kp1Bj_HH11_M23 and Kp2Bj_LN294_M23 were completely absorbed at 30 min. Kp10Bj_LN54_14 was not completely adsorbed at 60 min (Figure 6). One-step growth curve analysis of phages showed that all three phages have a latency time of 10 min. In addition, the burst sizes of phages Kp1Bj_HH11_M23, Kp2Bj_LN294_M23, and Kp10Bj_LN54_14 were 402, 2208, and 119 PFU/cell, respectively (Figure 7).

**Figure 5.**
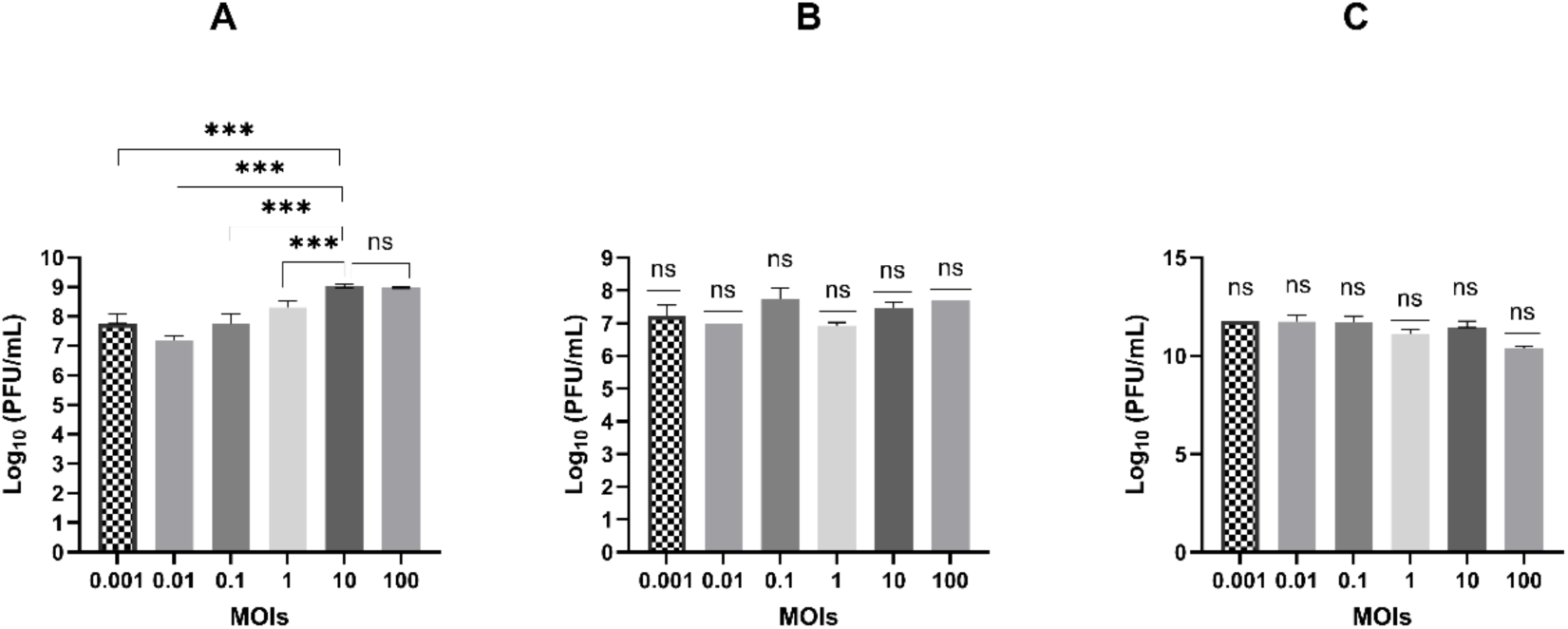
Titer of each phage at different MOIs. **A**: titer of Kp1Bj_HH11_M23 at different MOIs. **B**: titer of Kp2Bj_LN294_M23 at different MOIs. **C**: titer of Kp10Bj_LN54_14 at different MOIs. The data are presented as the mean ± standard deviation of three independent experiments. The asterisks indicate significant differences between the MOIs (*P<0.05, **P<0.01 or ***P<0.001, Ordinary one-way ANOVA followed by Tukey’s multiple comparisons test).

**Figure 6.**
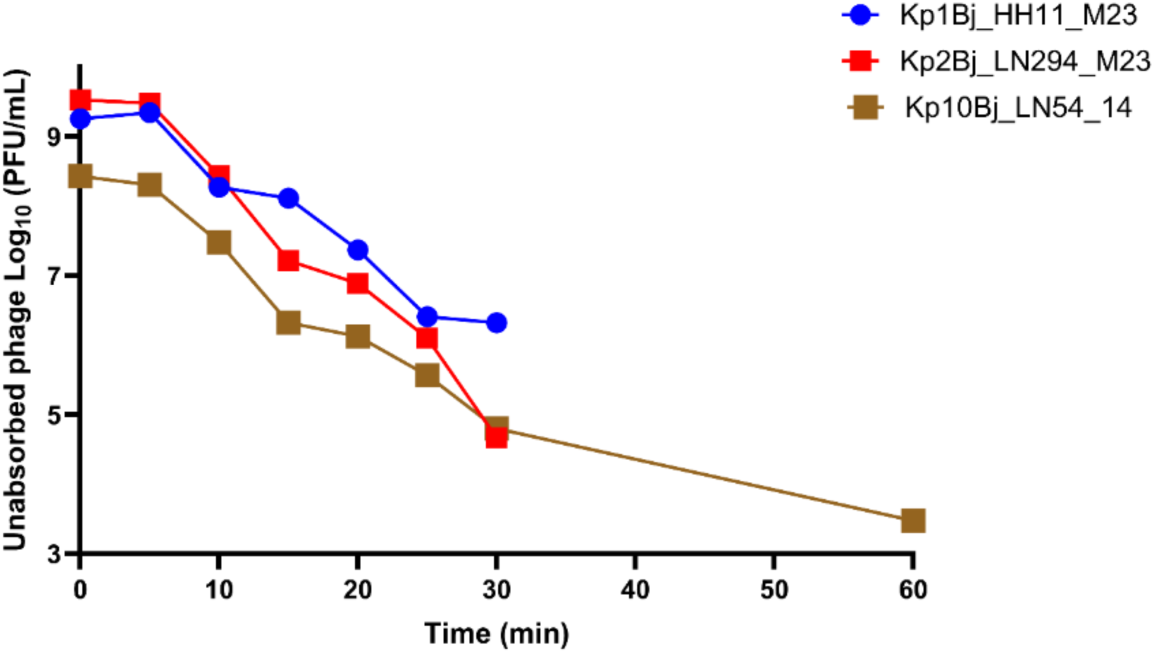
Kinetic adsorption of phages to the host strains surfaces at their optimal MOI.

**Figure 7.**
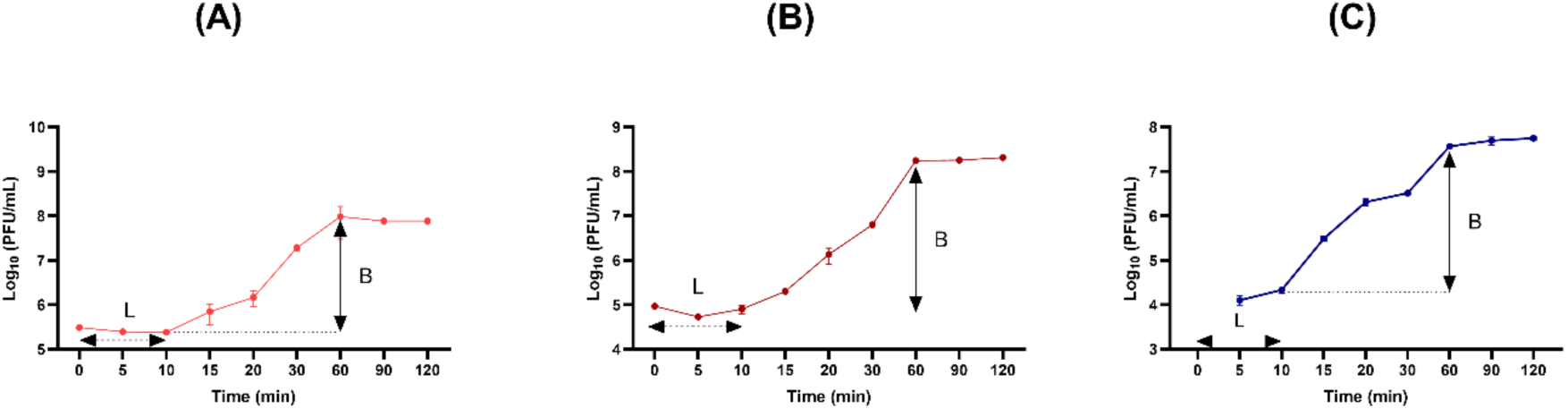
One-step growth curve analysis of phages at their optimal MOI. **(A)** one-step growth curve of Kp1Bj_HH11_M23. **(B)** one-step growth curve of Kp2Bj_LN294_M23. **(C)** one-step growth curve of Kp10Bj_LN54_14.

### 3.5 Antibiofilm activities

The phage-treated groups showed a reduction in the planktonic cells compared with the control group after both 24 h and 48 h of incubation. This decrease of planktonic cells by Kp1Bj_HH11_M23 was highly significant at 48 h and not significant at 24 h. Reduction of planktonic cells by Kp2Bj_LN294_M23 was significant in 24 h and very significant in 48 h. While the reduction of planktonic cells by phage Kp10Bj_LN54_14 was highly significant at 24 h (Figure 8A, 8B and 8C).

**Figure 8.**
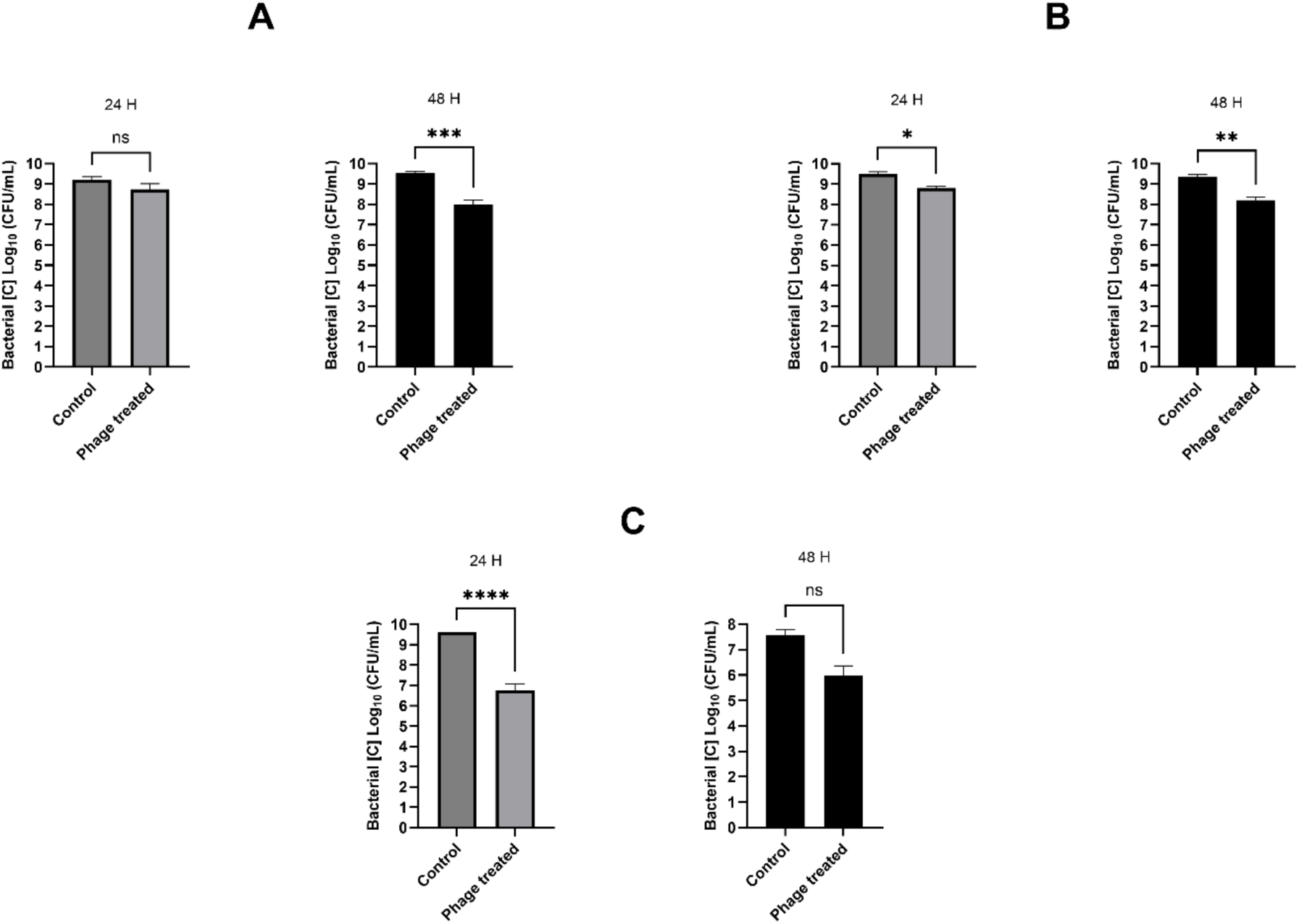
Impact of phage treatment on host cell viability, for 24 to 48 h before counting viable bacterial cells (CFU/ml). **A**: impact of Kp1Bj_HH11_M23 treatment on host cell viability. **B**: impact of Kp2Bj_LN294_M23 treatment on host cell viability. **C**: impact of Kp10Bj_LN54_14 treatment on host cell viability. The data are presented as the mean ± standard deviation of three independent experiments. The asterisks indicate significant differences between control and phage treated groups (*P<0.05, **P<0.01, ***P<0.001 or ****P<0.0001, unpaired t test).

Phages showed activity in destroying the biofilms formed within 24 and 48 hours. A decrease in the OD_600nm_ of biofilms in the phage-treated groups compared with the control groups was observed. However, the destructive effect of Kp1Bj_HH11_M23 and Kp10Bj_LN54_14 on the biofilms formed in 24 h was not significant and was very significant in 48 h compared to the control (Figures 9A and 9C). In contrast, the destruction of biofilm by Kp2Bj_LN294_M23 was highly significant at 24 h and not significant at 48 h compared to the control (Figure 9B).

**Figure 9.**
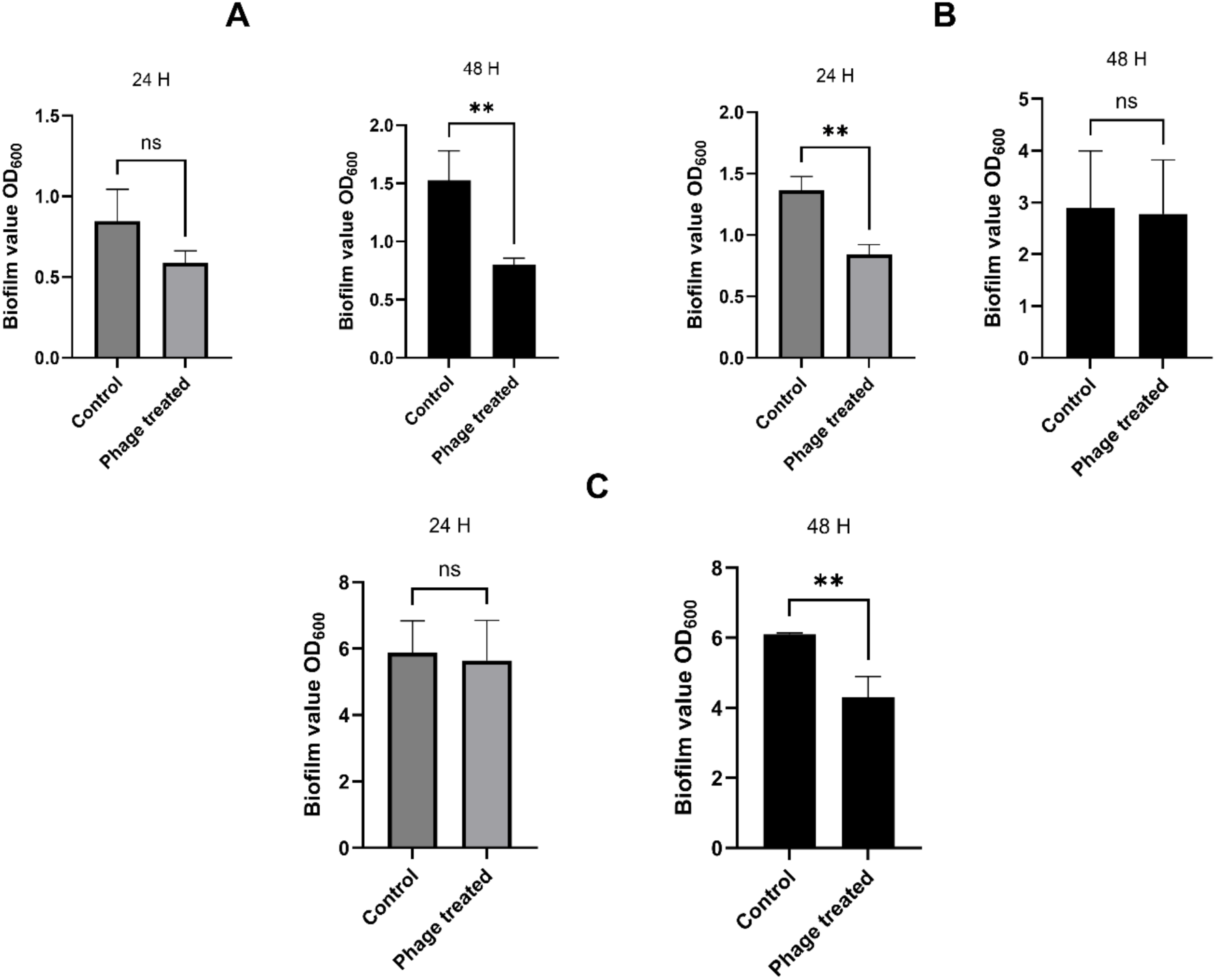
Antibiofilm activities of phages against host cells in LB medium. OD_600_ was measured after 24 and 48 h. **A**: antibiofilm activity of Kp1Bj_HH11_M23. **B**: antibiofilm activity of Kp2Bj_LN294_M23. **C**: antibiofilm activity of Kp10Bj_LN294_M67. The data are presented as the mean ± standard deviation of three independent experiments. The asterisks indicate significant differences between control and phage treated groups (*P<0.05, **P<0.01, ***P<0.001 or ****P<0.0001, unpaired t test).

### 3.6 Stability

Phages were sensitive to pH 1 and 2, losing their lytic activity on their respective hosts in an extremely acidic environment (Figures 10A, 10B, and 10C). Unlike Kp1Bj_HH11_M23 and Kp2Bj_LN294_M23, Kp10Bj LN54_14 was sensitive in pH 12, 13 and 14, and therefore inactive in an extremely basic environment (Figure 10C). Phages were stable in conditions with a pH between 3 and 11, inclusive, with titers from of 10^7^ to 10^9^, similar to their initial titers.

**Figure 10.**
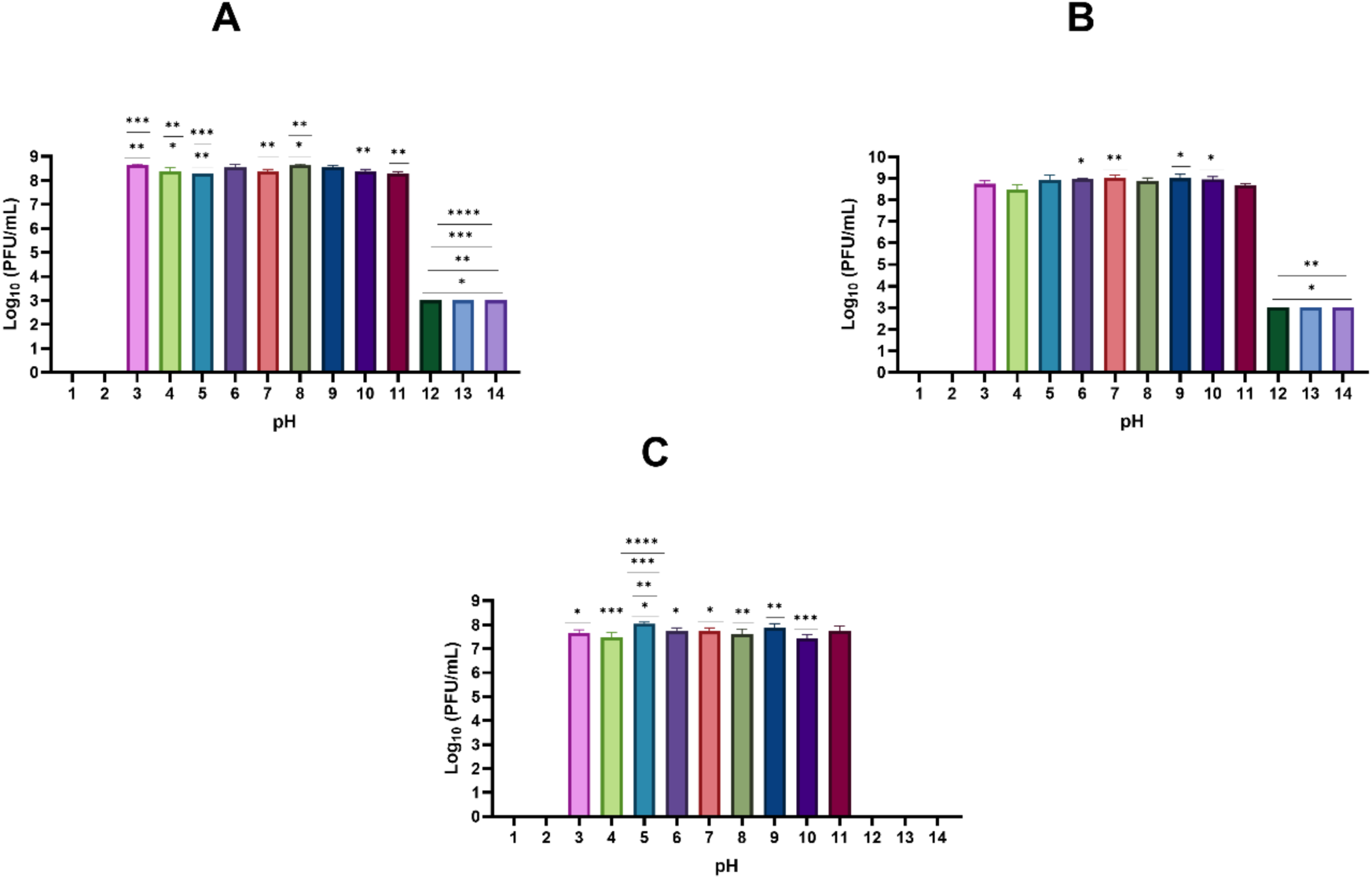
Stability of phages at different pH conditions (1 to 14). **A**: Kp1Bj_HH11_M23 stability. **B**: Kp2Bj_LN294_M23 stability. **C**: Kp10Bj_LN54_14 stability. The data are presented as the mean±standard deviation of three independent experiments. The asterisks indicate significant differences between the pH conditions (*P<0.05, **P<0.01, ***P<0.001 or ****P<0.0001, Ordinary one-way ANOVA followed by Tukey’s multiple comparisons test).

Above 70°C, phages showed no lytic activity on their respective hosts. Kp1Bj_HH11_M23 and Kp2Bj_LN294_M23 were stable over the temperature range from 4 to 60°C with a significant decrease in titers at 70°C (Figures 11A and 11B). Kp10Bj_LN54_14 was stable from 4 to 50°C and showed a significant decrease in titer at 60 and 70°C (Figure 11C).

**Figure 11.**
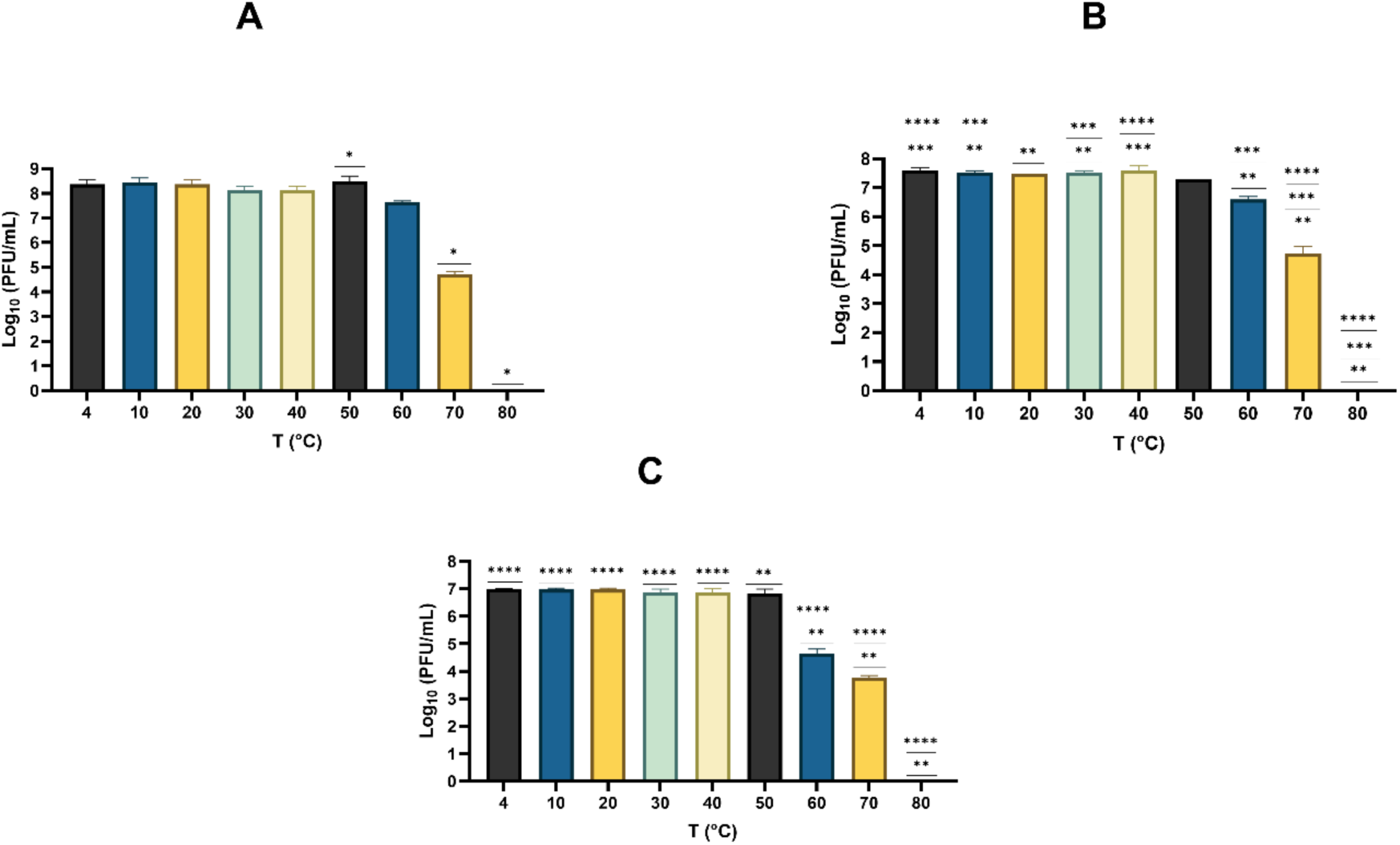
Stability of phages at different temperatures. **A**: Kp1Bj_HH11_M23 stability. **B**: Kp2Bj_LN294_M23 stability. **C**: Kp10Bj_LN54_14 stability. The data are presented as the mean ± standard deviation of three independent experiments. The asterisks indicate significant differences between temperature conditions (*P<0.05, **P<0.01, ***P<0.001 or ****P<0.0001, Ordinary one-way ANOVA followed by Tukey’s multiple comparisons test).

## 4 Discussion

The global increase in antimicrobial resistance (AMR) poses a significant challenge, particularly for low- and middle-income countries (LMICs), where roughly 90% of the worldwide deaths from AMR are expected to occur (Murray et al., 2022). This problem has led to the emergence of phage therapy as a promising alternative to antibiotics, whether used alone, in phage cocktails, or in combination with antibiotics. Lytic phages are considered one of the best solutions for treating infections caused by multidrug-resistant bacteria (Bleriot et al., 2023). Currently, most research on phage therapy is concentrated in high-income countries (Eskenazi et al., 2022; Lebeaux et al., 2021; Jennes et al., 2017; Leitner et al., 2017; Miedzybrodzki et al., 2012). There is no data available on the effectiveness of phage therapy in Africa, where bacterial epidemiology and healthcare infrastructure differ substantially from those in high-income countries. Furthermore, it is crucial to generate data on lytic phages that are candidates for therapeutic use in the treatment of infections caused by multidrug-resistant bacteria in Africa. The present study aims to evaluate the potential contributions of phages in reducing bacterial growth and degrading biofilms formed by clinical multidrug-resistant strains of *Klebsiella pneumoniae*. After characterizing the isolated phages in Benin, their antibacterial activity was evaluated *in vitro* in a biofilm model.

*K. pneumoniae* exhibits high resistance to a broad spectrum of drugs, including beta-lactams, fluoroquinolones and aminoglycosides, and is responsible for many human infections (Mulani et al., 2022). In this context, the use of specific and personalized phage cocktails may be essential in the fight against multidrug-resistant bacteria of the genus *Klebsiella* (Das and Kaledhonkar, 2024). For this, the constant isolation of novel *Klebsiella* phages capable of killing antibiotic-resistant *Klebsiella* and a thorough understanding of the biology of these phages are essential to develop new antibacterial strategies that could be used to treat human pathogens (Fayez et al., 2023). As part of this study, three novel Myoviruses were isolated in Benin from samples of hospital effluent (Kp1Bj_HH11_M23), stool (Kp10Bj_LN54_14), and animal feces (Kp2Bj_LN294_M23), respectively. Which shows that lytic phages can be found in human, animal and environmental samples. The titer of these phages is between 7.10^9^ to 2.10^10^ PFU/ml, implying that they have a high affinity for their host bacteria. These phages would have a capability to treat infection due to biofilm-producing bacteria such as *K. pneumoniae* (Horiuk et al., 2020). The three phages showed lytic activity on a wide range of clinical strains of *K. pneumoniae* (at least 40 strains) with various sequence types. Importantly, they can infect clinical isolates with different capsular loci, including KL114, KL53, KL8, KL62, KL64, KL124, KL102, KL57, KL39, KL48, KL17, KL20, KL2, KL30, KL51, KL105, KL146, KL28, KL122, KL38, KL3, and KL163. The diversity of *Klebsiella* capsules is the main obstacle to the development of broad-spectrum phage cocktails. Since many phages use specific capsular polysaccharides (CPS) as receptors, a single phage often infects only one or a few of the many known capsular serotypes of *Klebsiella*, which limits its clinical efficacy (Chen et al., 2024; Ferriol-González et al., 2024; Wang et al., 2024; Burke et al., 2025). These results show that the three phages are good candidates for development of broad-spectrum phage cocktails and therapy against infections (urinary tract infections, pneumonia, sepsis, nosocomial infections) caused by hypervirulent and antibiotic-resistant (even last resort) strains of *K. pneumoniae*. With a wide host range (from 68.97% to 74.14%), the three *Klebsiella* phages have a broader lytic spectrum than some previously characterized novel *Klebsiella* phages (Ciacci et al., 2018; Bleriot et al., 2023; Fang et al., 2023; Köhne et al., 2025), which lysed 30 to 40% (Köhne et al., 2025), 36.17 to 57.45% (Bleriot et al., 2023); 13.59% (Ciacci et al., 2018) and 2% (Fang et al., 2023) of the clinical isolates tested.

The morphology of the three phages (Kp1Bj_HH11_M23, Kp2Bj_LN294_M23 and Kp10Bj_LN54_14) was observed using a transmission electron microscope and indicated that they had Myovirus, and all belonged to the order Caudovirales. This result was confirmed by phylogeny, which showed the proximity of the three phages to other phages in the same family (Myoviridae). This classification corresponds to that of other phages known to infect clinical *K. pneumoniae* (Min et al., 2019; Zurabov and Zhilenkov, 2019; Fang and Zong, 2022; Fang et al., 2023; Gorodnichev et al., 2025; Gou et al., 2025). Furthermore, phages Kp1Bj_HH11_M23 and Kp2Bj_LN294_M23 belong to the Marfavirus genus, while Kp10Bj_LN54_14 belongs to the Jiaodavirus genus. Phylogeny has shown that phages Kp1Bj_HH11_M23 and Kp2Bj_LN294_M23 are closely related to phages Marfa (MN044033) and vB_Kpn_F48 (MG746602), which are also phages belonging to the genus Marfavirus (Ciacci et al., 2018; Harb et al., 2019). Similarly, the phylogenetic tree showed that Kp10Bj_LN54_14 is closely related to the phages PK0111 (NC_031095) and KPV15 (KY000080). PK0111 is a phage capable of infecting clinical strains of *K. oxytoca* and *K. pneumoniae* (Park et al., 2017). KPV15 is a phage that is part of the phage cocktail used for the treatment and prophylaxis of healthcare-associated infections in a mouse model and in phage therapy in patients with infections related to cardiothoracic surgery (Aleshkin et al., 2016; Rubalskii et al., 2020). Thus, Kp10Bj_LN54_14 could be a candidate for phage therapy against healthcare-associated infections caused by bacteria of the genus *Klebsiella*. Genome analysis showed that there are no genes associated with recombinases, integrases, or repressors in any of the three phage genomes, confirming that all three phages follow a lytic life cycle. The absence of genes linked to bacterial virulence or antibiotic resistance in each of the three phage genomes indicates that these new lytic phages meet the generally accepted genomic criteria for therapeutic phages (Al-Anany et al., 2021; Song et al., 2021).

The optimal multiplicity of infections (MOIs) of the three phages (Kp1Bj_HH11_M23, Kp2Bj_LN294_M23 and Kp10Bj_LN54_14) are 10, 0.1, and 0.001, respectively, where they produce the highest phage titers. The incubation time for each of the three phages is 10 minutes. According to Abedon (1989) and Chen et al. (2023), producing large quantities of phages using a lower MOI and a shorter incubation period may prove to be less expensive. Thus, phage Kp10Bj_LN54_14 may be considerably better than Kp1Bj_HH11_M23 and Kp2Bj_LN294_M23 in terms of optimal phage production. According to the one step growth curve, Kp2Bj_LN294_M23 has the largest burst size (2208 PFU/cell), followed by Kp1Bj_HH11_M23 (402 PFU/cell) and Kp10Bj_LN54_14 (119 PFU/cell). Likewise, Kp2Bj_LN294 is larger than previously reported *Klebsiella* phages vB_Kpn_ZC2 (650 PFU/cell), vB_KpnP_kP17 (331 PFU/cell), vB_KleS-HSE3 (277 PFU/cell) and IME183 (135 PFU/cell) (Xu et al., 2023; Abdel-Razek et al., 2025; Fayez et al., 2023; Peng et al., 2020). The rapid phage absorption and massive release of virions suggest that Kp1Bj_HH11_M23, Kp2Bj_LN294_M23 and Kp10Bj_LN54_14 might have robust therapeutic efficacy.

The time-killing assay enabled us to determine phage virulence against *K. pneumoniae* planktonic cells. The growth of planktonic cells was significantly inhibited by the three phages within 48 hours of incubation. Phages Kp1Bj_HH11_M23, Kp2Bj_LN294_M23 and Kp10Bj_LN54_14 demonstrated a significant ability to inhibit *Klebsiella* biofilm formation and degrade preformed biofilms in vitro. However, phage Kp2Bj_LN294_M23 showed an increase in biofilm formation 48 hours after treatment. The biofilm matrix of bacterial species generally consists of extracellular polymeric substances (polysaccharides, DNA, proteins, etc.) (Abdel-Razek et al., 2025; Wu et al., 2019). The increase in biofilm after 48 hours of treatment could be explained by the bacteria’s response to phage pressure, leading to the activation of defense mechanisms and thereby increasing the production of extracellular matrix, which strengthens the biofilm structure. Alternatively, the partial destruction of the biofilm by the phages may release extracellular substances that act as nutrients, promoting the adhesion and growth of a new, denser biofilm. These reactions highlight the importance of phage cocktails in biofilm treatment. Thus, the phage cocktail consisting of Kp1Bj_HH11_M23, Kp2Bj_LN294_M23, and Kp10Bj_LN54_14 could be a very good candidate for preclinical and clinical testing in the treatment of infections caused by hypervirulent, biofilm-producing, multidrug-resistant *K. pneumoniae* strains.

For therapeutic purposes, it is important to maintain phage viability under various stress conditions (Islam et al., 2023). According to Bai et al. (2022), the industrial application of phages involves exposing them to common conditions with a temperature range of 4 to 40°C and a pH ranging from 4 to 10 during production and processing. Kp1Bj_HH11_M23, Kp2Bj_LN294_M23, and Kp10Bj_LN54_14 remain stable within these temperature and pH ranges. All three phages retained partial activity under high-temperature conditions up to 70°C, indicating that they are thermally stable and can be stored effectively. The pH tolerance of Kp1Bj_HH11_M23, Kp2Bj_LN294_M23, and Kp10Bj_LN54_14 ranges from 3 to 11, indicating that they can survive in the intestine and adapt to the microenvironments of different infection sites.

## 5 Conclusion

In summary, this study characterized the first *Klebsiella pneumoniae* phages isolated in Benin, Kp1Bj_HH11_M23, Kp2Bj_LN294_M23, and Kp10Bj_LN54_14, which belong to the family *Myoviridae* and the order Caudovirales. Kp1Bj_HH11_M23 and Kp2Bj_LN294_M23 are *Marfavirus*, while Kp10Bj_LN54_14 is a *Jiaodavirus*. These three novel phages exhibit lytic activity against more than 40 virulent and multidrug resistant clinical strains of *K. pneumoniae*. The genomes of these phages do not contain resistance or virulence genes. Furthermore, they exhibit rapid absorption, short latency periods, and the ability to release hundreds or even thousands of virions per infected cell during growth, highlighting their therapeutic potential for treating severe infections caused by hypervirulent clinical strains of *K. pneumoniae*. Furthermore, *in vitro* antibiofilm tests demonstrated the efficacy of all three phages in inhibiting both the formation and degradation of preformed biofilms. The stability of these three phages under challenging temperature and pH conditions indicates that they are potential candidates for production and therapeutic application. However, our next step will be the production of phage cocktail using phages Kp1Bj_HH11_M23, Kp2Bj_LN294_M23, and Kp10Bj_LN54_14 and to evaluate the efficacy of this cocktail in an animal model infected with hypervirulent clinical strains of *K. pneumoniae*.

## 6 Conflict of Interest

The authors declare that the research was conducted in the absence of any commercial or financial relationships that could be construed as a potential conflict of interest.

## 7 Author Contributions

AJA conceptualized the study. AJA, KF, GRN, RB, SM and ED participated in sample collection, laboratory work, and data generation. AJA, YMGH and JBGH carried data analysis. AJA, YMGH, GRN and RB participated in the writing-original draft preparation. AJA, YMGH, VD, HB, LBM and MRJC participated in the writing-review and editing. MRJC supervised the study. AJA, YMGH, ED, and RN contributed to funding. All authors have read and agreed to the published version of the manuscript.

## 8 Funding

This work was supported by grants from Islamic Development Bank-The World Academy of Sciences (IsDB-TWAS) under the ID number 507179, and Africa Reseach Excellent Fund (AREF) under the reference AREF-312-AGBA-F-C0909. AJA as Iso Lomso Fellow was support by Stellenbosch Institute for Advanced Study (STIAS). AJA as G4 group leader was support by Institut Pasteur. The bioinformatics analyses conducted on computerome2 HPC was covered by YMGH under the MULTI-OMICS AAS/APTI/22/068 project, which received financial assistance of The Gates Foundation (Investment ID: INV-007175, formerly OPP1191735) through the African Postdoctoral Training Initiative (APTI) in partnership with the U.S. National Institutes of Health (NIH). Stellenbosch Institute for Advanced Study (STIAS) covering the article publishing charges (APCs) for this article.

## Acknowledgments

Thanks to Becky Mayer Centre for Phage Research, University of Leicester and Dr. Karen D. Adler for access bacteria strain collection. Thanks to Dr. Natalie Allcock from microscopy core facility of the University of Leicester, for support with TEM Imaging. Thanks to Marius Gnanvi, Rémi Sessou, and Dr. Arnaud Soha for support with sampling. We also thanks to Dr. Tobi Nagel for reviewing the manuscript.

## 9 Data Availability Statement

The sequences of phages Kp1Bj_HH11_M23, Kp2Bj_LN294_M23, and Kp10Bj_LN54_14 have been deposited in the GenBank under accession numbers PV784030, PV784031 and PV784032. The original contributions presented in the study are included in the article/Supplementary Material, further inquiries can be directed to the corresponding author.

## Notes

### Competing Interest Statement

The authors have declared no competing interest.

