## Supplement Table S1 for "Isolation and characterization of novel *Klebsiella* phages from Benin and their antibiofilm activities on multidrug resistant and hypervirulent strains of *Klebsiella pneumoniae*": Supplementary Table S1.docx

**Supplementary Table S1.** Multidrug-resistant hypervirulent strains of *Klebsiella pneumoniae* used in this study

| N° | Strains Lab ID | Location/source | Sequence type | O type | K type |
| --- | --- | --- | --- | --- | --- |
| 1 | KR312 | Leicester Royal Infirmary | ST147 | O2v1 | KL64 |
| 2 | KR358 | Leicester Royal Infirmary | ST336 | O5 | KL25 |
| 3 | KR431 | Leicester Royal Infirmary | ST37 | Unknown | KL118 |
| 4 | KR437 | Leicester Royal Infirmary | ST147 | O4 | KL15 |
| 5 | L02 | King’s College Hospital, London | ST219 | O1v1 | KL114 |
| 6 | L05 | King’s College Hospital, London | ST736 | O3/O3a | KL53 |
| 7 | L06 | King’s College Hospital, London | ST894 | O3b | KL58 |
| 8 | L10 | King’s College Hospital, London | ST215 | O1v1 | KL16 |
| 9 | L11 | King’s College Hospital, London | ST736 | 03/03a | KL53 |
| 10 | L13 | King’s College Hospital, London | ST1-1LV | O1v2 | KL8 |
| 11 | L15 | King’s College Hospital, London | ST736 | O3/3a | KL53 |
| 12 | L19 | King’s College Hospital, London | ST894 | O3b | KL58 |
| 13 | L21 | King’s College Hospital, London | ST736 | O3/O3a | KL53 |
| 14 | L30 | King’s College Hospital, London | ST441 | O2v1 | KL62 |
| 15 | L32 | King’s College Hospital, London | ST37 | OL103 | KL12 |
| 16 | M31 | Manchester Royal Infirmary | ST2096 | O1v1 | KL64 |
| 17 | M19 | Manchester Royal Infirmary | ST48 | O1v1 | KL124 |
| 18 | M23 | Manchester Royal Infirmary | ST48 | O1v1 | KL124 |
| 19 | M20 | Manchester Royal Infirmary | ST48 | O1v1 | KL124 |
| 20 | M34 | Manchester Royal Infirmary | ST307 | O2v2 | KL102 |
| 21 | M36 | Manchester Royal Infirmary | ST307 | O2v2 | KL102 |
| 22 | M39 | Manchester Royal Infirmary | ST307 | O2v2 | KL102 |
| 23 | M40 | Manchester Royal Infirmary | ST307 | O2v2 | KL102 |
| 24 | M41 | Manchester Royal Infirmary | ST307 | O2v2 | KL102 |
| 25 | M54 | Manchester Royal Infirmary | ST307 | O2v2 | KL102 |
| 26 | M61 | Manchester Royal Infirmary | ST307-1LV | O2v2 | KL102 |
| 27 | M63 | Manchester Royal Infirmary | ST13-1LV | O1v2 | KL57 |
| 28 | M71 | Manchester Royal Infirmary | ST13-1LV | O1v2 | KL57 |
| 29 | M45 | Manchester Royal Infirmary | ST985 | O1v2 | KL39 |
| 30 | M49 | Manchester Royal Infirmary | ST15 | O1v1 | KL48 |
| 31 | M57 | Manchester Royal Infirmary | ST1715 | O1v1 | KL17 |
| 32 | M13 | Manchester Royal Infirmary | ST268 | O2v1 | KL20 |
| 33 | M47 | Manchester Royal Infirmary | ST25 | O1v2 | KL2 |
| 34 | M53 | Manchester Royal Infirmary | ST234 | O1v2 | KL30 |
| 35 | M11 | Manchester Royal Infirmary | ST231 | O1v2 | KL51 |
| 36 | M17 | Manchester Royal Infirmary | ST11 | O2v2 | KL105 |
| 37 | M68 | Manchester Royal Infirmary | ST13 | O1v2 | KL64 |
| 38 | M74 | Manchester Royal Infirmary | ST48 | O1v1 | KL124 |
| 39 | M25 | Manchester Royal Infirmary | ST11 | O2v2 | KL105 |
| 40 | M28 | Manchester Royal Infirmary | ST784 | O1v2 | KL146 |
| 41 | M48 | Manchester Royal Infirmary | ST307 | O2v2 | KL102 |
| 42 | M32 | Manchester Royal Infirmary | ST870 | O1v1 | KL17 |
| 43 | M33 | Manchester Royal Infirmary | ST391 | O1v2 | KL30 |
| 44 | M37 | Manchester Royal Infirmary | ST20 | O1v2 | KL28 |
| 45 | M38 | Manchester Royal Infirmary | ST268 | O2v1 | KL20 |
| 46 | M42 | Manchester Royal Infirmary | ST1504-2LV | O5 | KL125 |
| 47 | M52 | Manchester Royal Infirmary | ST2141 | O2v1 | Unknown |
| 48 | M58 | Manchester Royal Infirmary | ST219 | O1v1 | KL114 |
| 49 | M65 | Manchester Royal Infirmary | ST29 | O1/O2v2 | Unknown |
| 50 | M06 | Manchester Royal Infirmary | ST12 | O2v2 | KL122 |
| 51 | M35 | Manchester Royal Infirmary | ST105-1LV | O2v2 | KL102 |
| 52 | M03 | Manchester Royal Infirmary | ST1074 | O2v2 | KL38 |
| 53 | M24 | Manchester Royal Infirmary | ST12 | O2v2 | KL122 |
| 54 | M69 | Manchester Royal Infirmary | ST394 | O2v2 | KL3 |
| 55 | M44 | Manchester Royal Infirmary | ST1451 | O4 | KL131 |
| 56 | M51 | Manchester Royal Infirmary | ST253 | O2v1 | KL163 |
| 57 | M67 | Manchester Royal Infirmary | ST17 | O5 | KL25 |
| 58 | M75 | Manchester Royal Infirmary | ST17 | O5 | KL25 |
| 59 | 9 | Suru-Léré’s Hospital, Benin | unknow | unknow | unknow |
| 60 | 14 | Suru-Léré’s Hospital, Benin | unknow | unknow | unknow |
