## Supplement Table S2 for "Isolation and characterization of novel *Klebsiella* phages from Benin and their antibiofilm activities on multidrug resistant and hypervirulent strains of *Klebsiella pneumoniae*": Supplementary Table S2.docx

**Supplementary Table S2.** Phage lab ID, samples from which they are isolated and GPS coordinates of sample collection site.

| **Phage ID** | **Samples code/samples** | **GPS Cordonates** | **Host bacteria^a^** |
| --- | --- | --- | --- |
| Kp1Bj_HH11_M23 | HH11/Hospital effluent | Mother & child hospital : 6°21'42.8''N 2°26'19.9° E 6.3618909, 2.4388730 9C6Q+QG5 Cotonou, Benin | M23 |
| Kp2Bj_LN294_M23 | LN294/Pig droppings | Pig farm: 1 6°27’56.9’’N 2°24’51.1’’E 6.4657950, 2.4142070 FC87+8M Ganvie, Benin | M23 |
| Kp10Bj_LN54_14 | LN54/Human stools | Pig farm :1 6°27’56.9’’N 2°24’51.1’’E 6.4657950, 2.4142070 FC87+8M Ganvie, Benin | 14 |

**^a^** supplementary Table S1
