## Supplement Table S3 for "Isolation and characterization of novel *Klebsiella* phages from Benin and their antibiofilm activities on multidrug resistant and hypervirulent strains of *Klebsiella pneumoniae*": Supplementary Table S3.docx

**Supplementary Table S3.** List of *Klebsiella pneumoniae* phage genomes used for phylogenetic analysis

| Phage | Family | GenBank accession no. | Genome size (bp) | Source | Reference |
| --- | --- | --- | --- | --- | --- |
| Kp1Bj_HH11_M23 | Myoviridae | PV784030 | 170568 | Hospital effluent | This study |
| Kp2Bj_LN294_M23 | Myoviridae | PV784031 | 170567 | Pig droppings | This study |
| Kp10Bj_LN54_14 | Myoviridae | PV784032 | 167615 | Human stools | This study |
| Mulock | Myoviridae | MN098327 | 43727 | Wastewater | Min et al. 2019 |
| JD001 | Myoviridae | NC_020204 | 48814 | Seawater | Cui et al. 2012 |
| vB_KpnS_FZ14 | Myoviridae | MK521906 | 49370 | Sewage | Zurabov and Zhilenkov, 2019 |
| vB_KpnM_KB57 | Myoviridae | NC_028659 | 142987 | Sewage | Volozhantsev et al. 2016 |
| vB_KpnM_BIS47 | Myoviridae | KY652726 | 147443 | Sewage plant | Lubudda et al. 2017 |
| ZCKP1 | Myoviridae | MH252123 | 150925 | Fresh water | Taha et al. 2018 |
| KPV15 | Myoviridae | KY000080 | 167034 | Wastewater | Aleshkin et al. 2016 |
| vB_KpnM_KpV477 | Myoviridae | NC_031087 | 168272 | Clinical sample | Komisarova et al. 2017 |
| Marfa | Myoviridae | MN044033 | 168532 | Swine faeces | Harb et al. 2019 |
| PKO111 | Myoviridae | NC_031095 | 168758 | Sewage | Park et al. 2017 |
| vB_Kpn_F48 | Myoviridae | MG746602 | 170764 | Sewage | Ciacci et al. 2018 |
| KP27 | Myoviridae | NC_080080 | 174413 | Wastewater plant | Kesik-Szeloch et al. 2013 |
| KP15 | Myoviridae | NC_014036 | 174436 | Irrigated fields | Kesik-Szeloch et al. 2013 |
| PMBT1 | Myoviridae | LT607758 | 175206 | Sewage | Koberg et al. 2017 |
| Miro | Myoviridae | KT001919 | 176055 | Sewage | Mijalis et al. 2015 |
| Matisse | Myoviridae | NC_028750 | 176081 | Sewage | Provasek et al. 2015 |
| Patroon | Podoviridae | MK608335 | 39442 | Wastewater plant | Tran et al. 2019 |
| vB_KpnS_FZ12 | Podoviridae | MK521905 | 39519 | Sewage | Zurabov and Zhilenkov, 2019 |
| vB_Kp1 | Podoviridae | NC_028688 | 40114 | Wastewater plant | Alvez et al. 2029 |
| K5-4 | Podoviridae | KY389316 | 40163 | Sewage | Hsieh et al. 2017 |
| Henu1 | Podoviridae | MK203841.1 | 40352 | Sewage | Teng et al. 2019 |
| vB_KpnP_KpV767 | Podoviridae | KX712070 | 40395 | Sewage | Solovieva et al. 2018 |
| vB_KpnP_PRA33 | Podoviridae | KY652723 | 40605 | Sewage plant | Lubudda et al. 2017 |
| vB_KpnP_KpV763 | Podoviridae | KX591654 | 40765 | Sewage | Solovieva et al. 2018 |
| vB_KpnP_KpV289 | Podoviridae | NC_028977 | 41054 | Untreated sewage | Volozhantsev et al. 2016 |
| vB_KpnP_KpV766 | Podoviridae | KX712071 | 41283 | Sewage | Solovieva et al. 2018 |
| KPP5665-2 | Siphoviridae | MF695815 | 39241 | Mastitis milk | Carl et al. 2017 |
| Sushi | Siphoviridae | NC_028774 | 48754 | Sewage | Nguyen et al. 2015 |
| Sanco | Siphoviridae | MK618657 | 48790 | Wastewater plant | Richardson et al. 2019 |
| KLPN1 | Siphoviridae | NC_028760 | 49037 | Human caecum | Hoyles et al. 2015 |
| Shelby | Siphoviridae | MK931445 | 49045 | Pond water | Saldana et al. 2019 |
| vB_KpnS_GH-K3 | Siphoviridae | MH844531.1 | 49427 | Sewage | Gu et al. 2012 ; Cai et al. 2019 |
| 1513 | Siphoviridae | NC_028786 | 49462 | Sewage | Cao et al. 2015 |
| MezzoGao | Siphoviridae | MF612072 | 49807 | Wastewater plant | Gao et al. 2017 |
| KP36 | Siphoviridae | NC_029096 | 49818 | Wastewater plant | Kesik-Szeloch et al. 2013 |
| TSK1 | Siphoviridae | MH688453 | 49861 | Sewage | Tabassum et al. 2018 |
| Sin4 | Siphoviridae | MK931442 | 49916 | Wastewater plant | Castillo et al. 2019 |
| Skenny | Siphoviridae | MK931444 | 49935 | Activated sludge | Gramer et al. 2019 |
| vB_KpnS_FZ10 | Siphoviridae | MK521904 | 50381 | Sewage | Zurabov and Zhilenkov, 2019 |
